# Loss of PGC1α drives extracellular matrix remodelling in prostate cancer through CTHRC1

**DOI:** 10.64898/2026.05.05.722659

**Authors:** A Gonzalo-Paulano, A Macchia, A Schaub-Clerigué, B Lectez, J Gardoki, I Lejona, U Armendariz-Martínez, N Martín-Martín, I Astobiza, P Alonso, M Azkargorta, F Elortza, AM Aransay, I Rondon-Lorefice, I Mendizabal, L Egia-Mendikute, A Palazon, I Hermanova, A Azueta, A Loizaga, A Ugalde-Olano, A Carracedo, JJ Bravo-Cordero, L Valcarcel-Jimenez, V Torrano

## Abstract

Despite the high curation rate of localized prostate cancer, the fraction of patients that progress to metastasis still accounts for thousands of deaths worldwide, underscoring the need to identify early molecular events that prime tumours for aggressive disease. Here, we demonstrate that loss of the metabolic transcriptional coactivator PGC1α drives early extracellular matrix (ECM) remodelling in PCa, functionally linking epithelial transcriptional programs to tumour–microenvironment interactions.

Using genetically engineered mouse models, we show that combined deletion of *Pten* and *Pgc1α* induces early activation of ECM-related transcriptional programs, increased collagen deposition, and a transition towards an aligned collagen fibre architecture—hallmarks of aggressive disease—prior to metastatic dissemination. Consistently, human prostate tumours with low PGC1α expression display increased collagen deposition, supporting the clinical relevance of these findings. Restoration of PGC1α expression in prostate cancer cells suppresses cell adhesion to multiple ECM substrates, disrupts collagen organization, and impairs tumour growth in a transcription-dependent manner.

Through integrative matrisome proteomics and transcriptomics, we identify the secreted glycoprotein CTHRC1 as a key downstream effector that enhances the prognostic value of PGC1α in PCa patients. Functional loss- and gain-of-function studies establish CTHRC1 expression as both necessary and sufficient to restore ECM adhesion, cytoskeletal organization, collagen architecture, and tumorigenic capacity in PGC1α-expressing cells. Importantly, recombinant CTHRC1 rescues adhesion defects, indicating that its extracellular pool mediates this phenotype, whereas deglycosylation abolishes its pro-adhesive function, revealing a mechanistic requirement for glycosylation.

Collectively, our findings uncover an early, cell-intrinsic ECM remodelling program driven by PGC1α loss and identify the PGC1α–CTHRC1 axis as a mechanistic and clinically relevant regulator of PCa aggressiveness.

## INTRODUCTION

Prostate cancer (PCa) is the second most frequently diagnosed malignancy in men and the fifth leading cause of cancer-related mortality worldwide (https://gco.iarc.fr/today/home and (1)). While most patients are initially diagnosed with localized disease and respond well to conventional therapies (2), approximately 10% of PCas will eventually progress to an aggressive, metastatic stage that remains largely incurable and accounts for most PCa–related deaths (3). A major clinical challenge in PCa management is the lack of robust prognostic tools capable of identifying, at the time of diagnosis (4), those tumours harbouring an intrinsic potential for aggressive progression. This limitation reflects an incomplete understanding of the early molecular events that precede metastatic dissemination and underscores the need to define molecular and biological programs that prime prostate tumours for malignancy before overt metastasis becomes a clinical problem.

We and others previously addressed this challenge by integrating patient-derived datasets, genetically engineered mouse models, and *in vitro* experimental systems, leading to the identification of the transcriptional coactivator PGC1α as a tumour and metastasis suppressor in prostate cancer (5–9). However, these observations were largely derived from analyses of advanced-stage disease, limiting insight into the early biological processes triggered by PGC1α loss that may predispose prostate tumours to aggressive behaviour.

Beyond cell-intrinsic oncogenic alterations, increased evidence indicates that tumour progression is critically influenced by reciprocal interactions between malignant epithelial cells and the extracellular matrix (ECM) (10). Several cancer-derived ECM-associated proteins have been implicated in collagen organization and cell–matrix interactions in cancer (11–14) however, their regulation by epithelial cell-intrinsic transcriptional programs in PCa remains poorly defined.

The ECM is recognized as a dynamic and bioactive component of the tumour microenvironment that regulates cell adhesion, polarity, migration, and signal transduction (15). Alterations in ECM composition and organization—particularly changes in collagen deposition and fibre alignment—have been shown to promote invasive behaviour and metastatic competence across multiple cancer types (16). In this context, the matrisome, defined as the ensemble of ECM and ECM-associated proteins, has emerged as a key mediator of tumour–microenvironment crosstalk (17). Importantly, remodelling of the matrisome can occur early during tumour evolution and actively shape disease trajectory (18). Despite these advances, the contribution of early ECM remodelling to PCa progression, as well as the epithelial transcriptional programs that initiate these changes, remain poorly understood.

PGC1α is uniquely positioned to regulate these processes, as beyond its canonical role in metabolism, it functions as a pleiotropic transcriptional coactivator that drives transcriptional rewiring programs influencing cytoskeletal organization, cell-extrinsic signalling, and tumour–microenvironment interactions in prostate cancer (5, 6, 9). These observations led us to hypothesize that loss of PGC1α in prostate epithelial cells may initiate early alterations in cell–stroma interactions, thereby priming tumours for aggressive progression.

In this study, we combine genetically engineered mouse models, PCa cell lines, *in vivo* and *in ovo* tumour models, multi-omics analyses, and patient-derived transcriptomic datasets to investigate the consequences of PGC1α alteration on ECM biology in PCa. We demonstrate that PGC1α deficiency drives early transcriptional and structural remodelling of the ECM, leading to altered cell–matrix interactions and collagen organization associated with aggressive disease. Through integrated transcriptomic and matrisome-specific proteomic approaches, we identify a PGC1α-CTHRC1 ECM regulatory axis that controls these phenotypes and predicts clinical outcome in PCa patients. Collectively, our findings uncover an early cell-intrinsic driven mechanism linking metabolic regulation, ECM remodelling, and PCa aggressiveness.

## RESULTS

### Loss of PGC1α drives early ECM remodelling in prostate cancer

We sought to characterize the early oncogenic events that precede the metastatic phenotype associated with the loss of PTEN and PGC1α in prostate epithelial cells (8), with the aim of identifying alterations linked to disease aggressiveness. To this end, we performed a pathological analysis of prostate tumours derived from 3-month-old *Pten* knockout (*Pten^pc-/-^ Pgc1a^pc+/+^*) and *Pten/Pgc1α* double-knockout (*Pten^pc-/-^Pgc1a^pc-/-^*) mice. Consistent with previous reports (19), prostate-specific deletion of *Pten* at this early time point resulted predominantly in high-grade prostatic intraepithelial neoplasia (HGPIN) lesions, with a low incidence of carcinoma (Figure 1 A). In contrast, combined loss of *Pten* and *Pgc1α* significantly increased the incidence of invasive carcinoma lesions (Figure 1 A, Supplementary Figure 1 A), indicating enhanced tumour aggressiveness at early stages. Importantly, these differences were not accompanied by changes in tumour weight (anterior prostate), stromal fibrosis, immune infiltration, or detectable metastatic dissemination (Supplementary Figure 1 B–E), revealing that PGC1α loss promotes early tumorigenesis independently of overt stromal or metastatic alterations.

**Figure 1.**
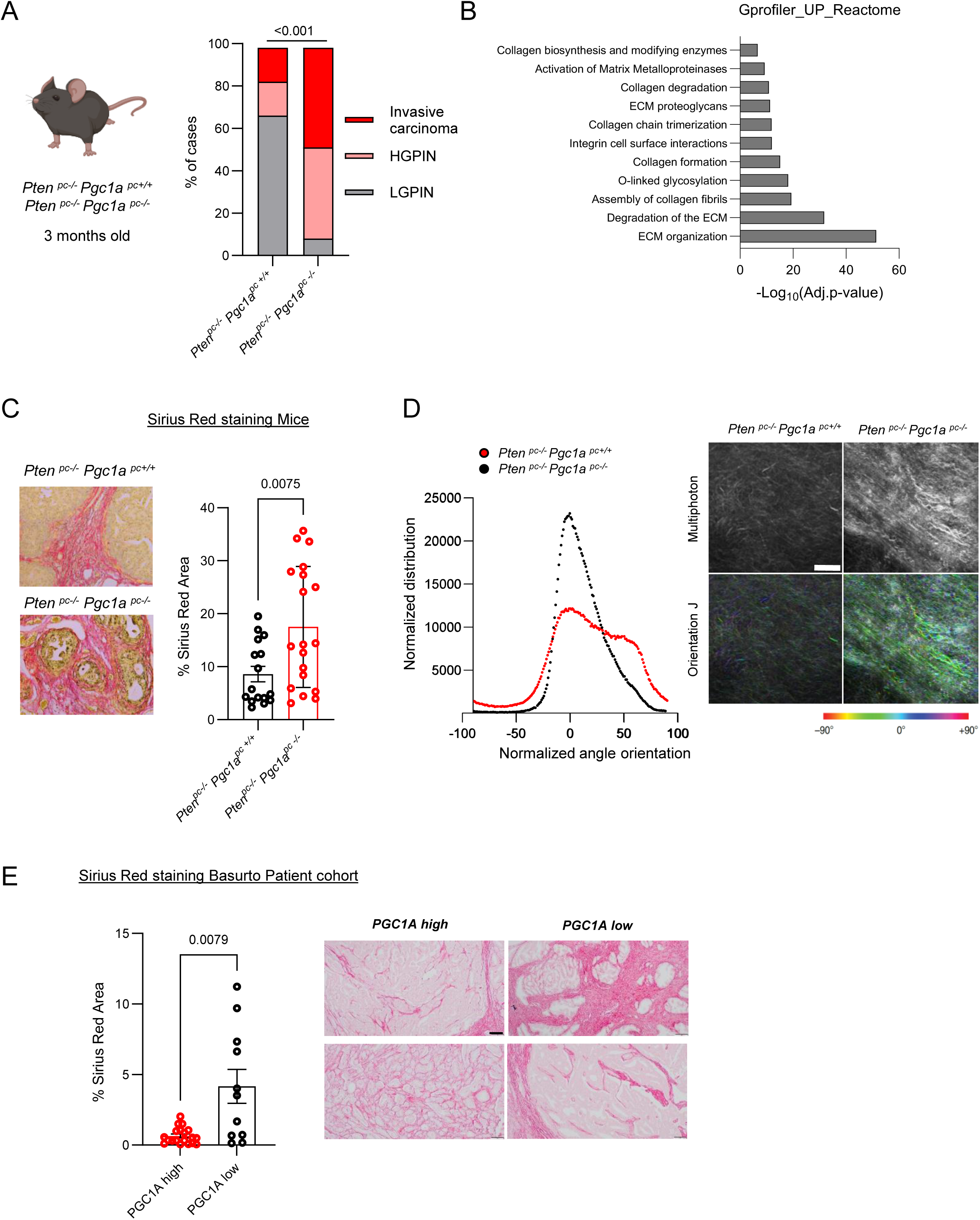
Double deletion of *Pten* and *Pgc1α* induces ECM remodelling in prostate cancer. **A**. Histopathological analysis performed in hematoxylin-eosin stained prostate tumours harvested from *Pten^pc-/-^ Pgc1a^p+/+^* and *Pten^pc-/-^ Pgc1a^pc-/-^* 3-months old mice (Mice sample size *Pten^pc-/-^ Pgc1a^p+/+^* n=12 and *Pten^pc-/-^ Pgc1a^pc-/-^* n=23). **B**. Functional enrichment analysis of DEG genes whose expression was increased in *Pten^pc-/-^ Pgc1a^pc-/-^* prostate tumours compared to *Pten^pc-/-^ Pgc1a^p+/+^*. **C**. Quantification of collagen content in 3-months old *Pten^pc-/-^ Pgc1a^p+/+^* and *Pten^pc-/-^ Pgc1a^pc-/-^* prostate tumours stained with Picros Sirius Red (Image sample size *Pten^pc-/-^ Pgc1a^p+/+^* n= 16 images/areas (3 tumours) and *Pten^pc-/-^ Pgc1a^pc-/-^* n= 16 images/areas (4 tumours). **D**. Analysis of collagen fibre orientation in decellularized 3-months old *Pten^pc-/-^ Pgc1a^p+/+^* and *Pten^pc-/-^ Pgc1a^pc-/-^*luciferase prostate tumours using multiphoton microscopy (n=2 x genotype). Scale bars represent 100 μm and colour coding represents angle degree. **E**. Quantification of collagen content in PCa patients (Rondon *et al*) stratified according to PGC1a mRNA expression (n=12 patients/30 images). HGPIN: high grade prostatic intraepithelial neoplasia. LGPIN: low grade prostatic intraepithelial neoplasia. UP: upregulated. Statistics: Fisheŕs exact test (A), unpaired t-test (C, E). Numbers represent p-values. Error bars indicate s.e.m.

To investigate the molecular basis of this phenotype, we performed bulk RNA-sequencing of *Pten^pc-/-^ Pgc1a^pc+/+^* and *Pten^pc-/-^ Pgc1a^pc-/-^* tumours. Differential expression analysis revealed extensive transcriptional reprogramming upon Pgc1α deletion, including 1440 upregulated and 874 downregulated protein-coding genes (Supplementary Figure 1 F and Table S1). Notably, beyond metabolic pathway alterations, these were significantly enriched for gene signatures related to ECM organization, collagen biosynthesis and degradation, and cell–matrix interactions (Figure 1 B), pointing to ECM production and remodelling as a prominent feature of PGC1α-deficient tumours.

Consistent with these transcriptional alterations, histological analysis revealed increased collagen deposition in *Pten^pc-/-^ Pgc1a^pc-/-^* tumours, as quantified by picrosirius red staining (Figure 1 C), and no significant changes in immune cell populations (Supplementary Figure 1 H). These observations prompted us to further analyse the ECM in these tumours.

Given that fibroblasts and cancer-associated fibroblasts (CAFs) are the primary producers of ECM components, we next assessed their potential contribution to this phenotype. Projection of established prostate CAF gene signatures (20, 21) onto our RNA-seq data revealed no significant differences between *Pten^pc-/-^ Pgc1a^pc+/+^* and *Pten^pc/-^ Pgc1a^pc-/-^* tumours (Supplementary Figure 1 I). These findings show that the ECM-related transcriptional program induced by PGC1α loss is not associated to molecular changes characteristic of the stromal compartment but may originate from epithelial tumour cells.

In addition to changes in ECM composition, alterations in collagen architecture are key determinants of tumour aggressiveness. While normal tissues exhibit curved, isotropic collagen fibres, malignant lesions are characterized by aligned, anisotropic structures (22). Multiphoton microscopy revealed that *Pten^pc-/-^ Pgc1a^pc+/+^* tumours displayed predominantly isotropic collagen organization, whereas *Pten^pc-/-^ Pgc1a^pc-/-^* tumours exhibited highly aligned collagen fibres (Figure 1 D), consistent with a more aggressive phenotype.

To assess the clinical relevance of these observations, we analysed a cohort of human primary prostate tumours with matched transcriptomic and histological data (23). Consistent with previous reports, low PGC1α expression was associated with poor prognosis (Supplementary Figure 1 J). Importantly, tumours with low PGC1α levels displayed increased collagen deposition compared to those with higher expression (Figure 1 E), recapitulating our observations in mouse models.

Collectively, these data indicate that PGC1α loss in the mouse prostate epithelium drives early ECM remodelling, characterized by increased collagen deposition and fibre alignment, independently of stromal activation, and represents a hallmark of aggressive PCa.

### PGC1α suppresses cell–ECM adhesion and ECM organization through ERRα-dependent transcriptional control

Given the ECM remodelling phenotype observed upon PGC1α loss *in vivo*, we next investigated whether PGC1α regulates cell–ECM interactions. To this end, we employed a doxycycline-inducible system that enables the restoration of PGC1α expression (Supplementary Figure 2 A), which elicits a tumour-suppressive phenotype in PCa cancer cell lines (8). RNA sequencing analysis of PC3 cells with or without PGC1α expression (Supplementary Table S4) confirmed activation of metabolic programs and the repression of cell cycle–associated pathways, as previously described (Supplementary Figure 2 B-C). Notably, PGC1α re-expression also led to a marked downregulation of genes involved in cell–ECM interactions and ECM organization (Figure 2 A), mirroring our *in vivo* observations.

**Figure 2.**
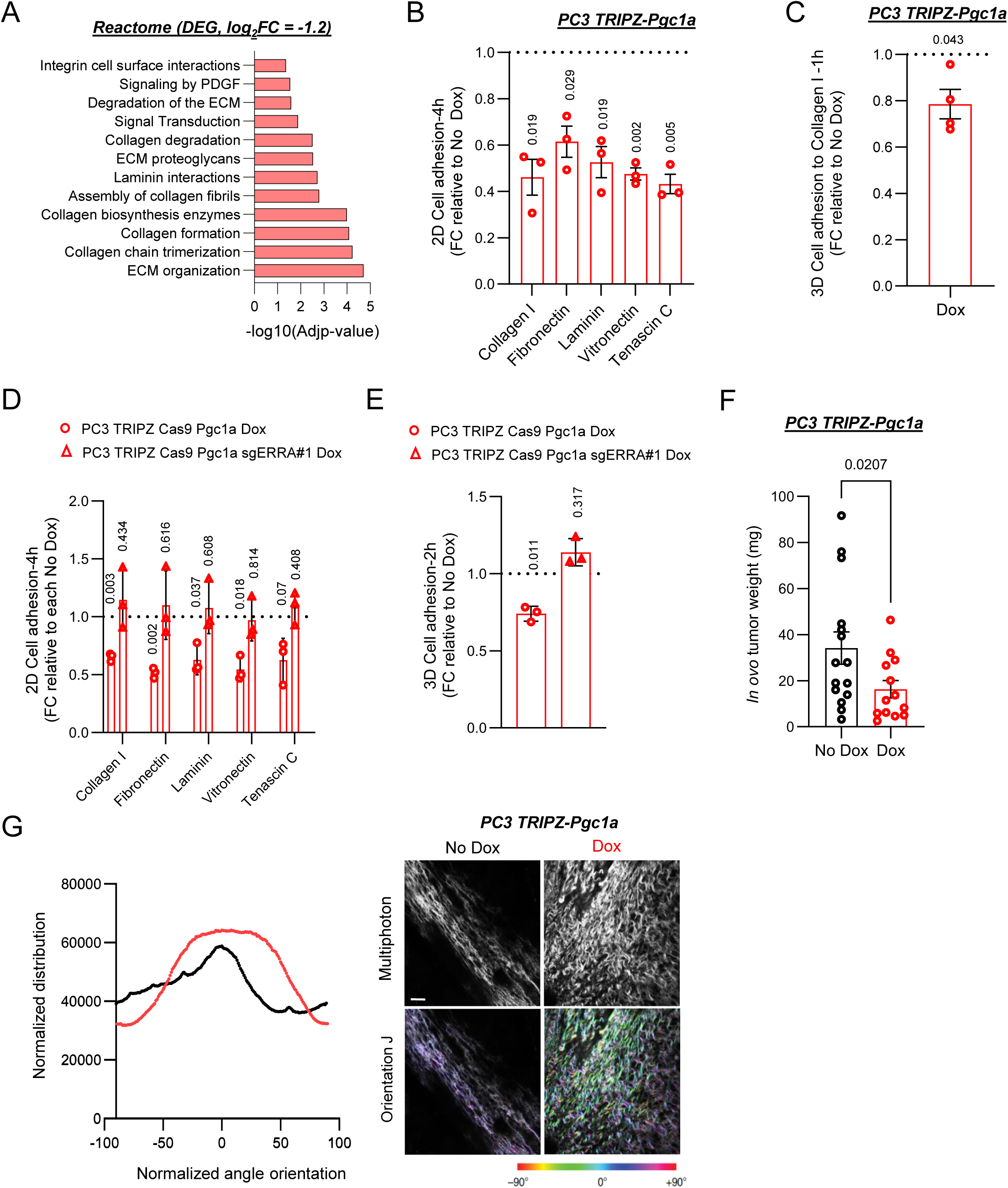
PGC1α reduces prostate cancer cell-ECM adhesion and remodels ECM organisation. **A.** Functional enrichment analysis of genes differentially downregulated in PC3 cells upon PGC1α re-expression. **B**. Effect of PGC1α re-expression in the capacity of PC3 cells to adhere to a panel of 2D extracellular matrixes *in vitro* (n=3). **C**. Effect of PGC1α re-expression in the capacity of PC3 cells to adhere to 3D collagen matrix *in vitro* (n=4). **D-E**. Effect of ERRα deletion in the PGC1α-driven phenotype on 2D (D) coating and 3D-collagen I matrix (C) cell-adhesion (n=3). **F**. Effect of PGC1α re-expression in the capacity of PC3 to form tumours *in ovo* (n=13-15 tumours). **G**. Quantification of collagen fibre alignment (left panel) in tumours generated *in ovo* by PGC1a-expressing and non-expressing PC3 cells. Right panel correspond to representative second harmonic generation multiphoton microscopy images of the aforementioned *in ovo* tumours. Scale bar represents 20µm and colour coding represents angle degree. In panels B to E, data are normalized to the -Dox (non-PGC1α expressing) conditions, depicted by a black dotted line. DEG: differential expressed genes; FC: fold change; Dox: doxycycline. Statistics: one sample t-test with reference value 1 (B, C, D and E), unpaired t-test (F). Numbers represent p-values. Error bars indicate s.e.m.

We next asked whether these transcriptional changes translate into functional alterations in cell–matrix interactions. Strikingly, PGC1α re-expression significantly decreased the ability of PC3 cells to adhere to multiple ECM substrates in two-dimensional (2D) condition, with the strongest effect observed on collagen I, and in three-dimensional (3D) collagen matrixes (Figure 2 B–C). This phenotype was reproducible in an independent PCa cell line (DU145; Supplementary Figure 2 D–E) and was not affected by doxycycline treatment (Supplementary Figure 2 F–G.

Importantly, genetic ablation of ERRα (9), a key transcriptional partner of PGC1α, fully rescued the adhesive capacity of PGC1α-expressing cells in both 2D and 3D conditions (Figure 2 D–E). These results reveal an unprecedented regulation of cell–ECM adhesion by PGC1α that is strictly dependent on its cooperation with ERRα.

We next examined whether PGC1α also influences the ability of tumour cells to modulate the ECM *in vivo*. Using an *in ovo* tumour model, we validated that PGC1α re-expression markedly reduced tumour growth (Figure 2 F), consistent with its tumour-suppressive role (6, 8, 9). Notably, collagen fibres within PGC1α-expressing tumours displayed a disorganized and isotropic architecture (Figure 2 G, lower panel), feature of indolent lesions (12). In contrast, PGC1α non-expressing tumours exhibited aligned collagen fibres (Figure 2 G, upper panel), associated with aggressive tumour phenotypes (12).

Collectively, these findings demonstrate that PGC1α-ERRα restrains epithelial cell–ECM adhesion and ECM organization, thereby limiting the acquisition of pro-tumourigenic ECM features.

### PGC1α-dependent matrisome remodelling identifies CTHRC1 as a candidate downstream effector

To ascertain the molecular mediators of the effect of PGC1α on cell–ECM interactions and ECM organization , we analysed the tumour matrisome (24) using an PCa model system in which tumour cells are implanted in their natural site through orthotopic administration.

PGC1α-positive and negative PC3 cells were orthotopically injected in the ventral lobe of the prostate of nude mice, and the matrisome composition of the resulting tumours was characterised. Consistent with the established tumour-suppressive role of PGC1α in PCa, re-expression of PGC1α in PC3 cells (Supplementary Figure 3 A) resulted in reduced tumour growth and metastatic dissemination to the lymph nodes (8) (Figure 3 A and Supplementary Figure 3 B). Label-free quantitative proteomic analysis (Supplementary Figure 3 C and Table S5) revealed extensive remodelling of the tumour-associated proteome, with a substantial fraction of differentially detected proteins corresponding to human matrisome components (Figure 3 B). Since we observed almost an absence and no differences in the stromal fraction by histological analysis, we hypothesized that changes in the abundance of matrisome-associated proteinswere due to tumour cell–intrinsic alterations.

**Figure 3.**
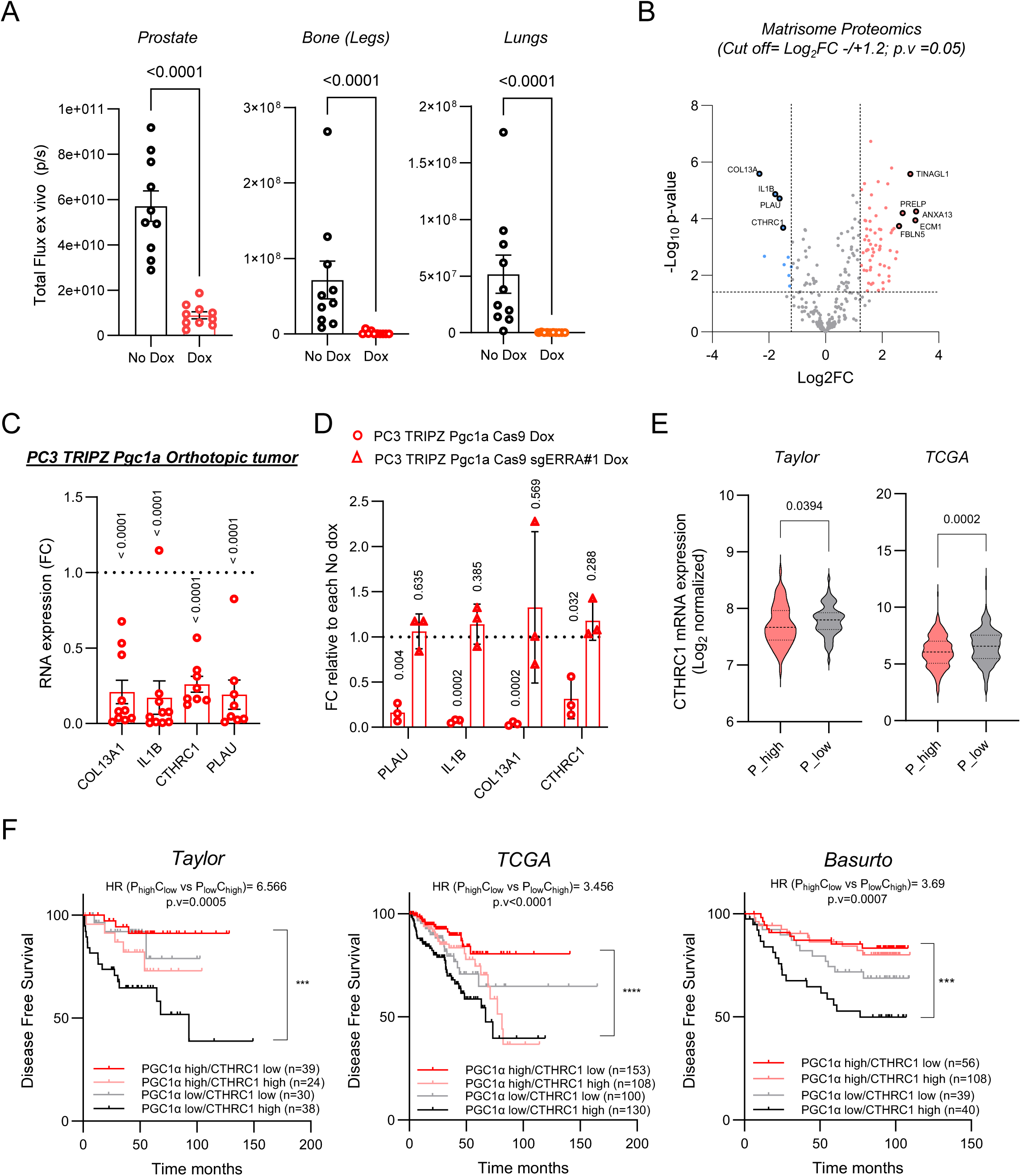
PGC1α-dependent matrisome modulation Identifies CTHRC1 as a candidate effector. **A.** Effect of PGC1α re-expression in the capacity of PC3 cells to form and grow orthotopic prostate tumours (left panel) and to metastazise to the bone and lungs (right panels). PC3-luc expressing and non-expressing cells were quantified by *in vivo* imaging of the prostate and *ex vivo* imaging of bone legs and lungs (n=10). **B**. Volcano plot representing label-free LC/MS data of matrisome-related human proteins differentially detected in orthotopic tumours generated by PGC1α-expressing tumours compared to non-expressing ones. **C**. Gene expression analysis (RT-qPCR) of candidate matrisome-related genes in PGC1α expressing orthotopic tumours compared to the negative ones (n=8-10). Data was normalized to the average mRNA expression of each gene in PGC1α-non expressing tumours, represented as a dotted line. **D**. Effect of ERRα deletion in the PGC1α-driven transcriptional downregulation of candidate matrisome-genes (RT-qPCR, n=3) *in vitro* (PC3 cells). Data was normalized to each No Dox condition, depicted as a dotted line. **E**. Analysis of CTHRC1 mRNA in PCa patients stratified according to the mean expression of PGC1α mRNA (PPARGC1A). Dox: doxycycline; FC: fold change; P: PGC1α. Sample sizes: Taylor database n=131 and TCGA provisional, n=497. Statistics: Unpaired t-test (A), one sample t-test with reference value 1 (C and D) and Mann Whitney test (E). Numbers represent p-values. Error bars indicate s.e.m.

To identify candidate mediators of the PGC1α-driven phenotype, we integrated transcriptomic (Figure 2 A) and matrisome proteomic data (Figure 3 B). This analysis revealed four matrisome-associated candidates—COL13A1, CTHRC1, IL1B, and PLAU—that were downregulated upon PGC1α re-expression (Supplementary Figure 3 D). The transcriptional regulation of these genes by PGC1α was validated both in orthotopic tumours (Figure 3 C) and *in vitro* across independent PCa cell lines (Figure 3 D and Supplementary Figure 3 E). In line with our previous findings, PGC1α-driven transcriptional regulation of the four candidates was dependent on ERRα (Figure 3 D).

Among these 4 genes, CTHRC1 emerged as the only one exhibiting a robust and consistent inverse correlation with PGC1α expression across independent human prostate cancer cohorts (Figure 3 E and Supplementary Figure 3 F). Importantly, analysis of laser-microdissected tumour samples (25) revealed that this inverse relationship was restricted to epithelial compartments and was not observed in the stroma (Supplementary Figure 3 G), supporting a tumour cell–intrinsic regulation.

### The PGC1α–CTHRC1 transcriptional axis predicts clinical outcome in prostate cancer

Given the inverse relationship between PGC1α and CTHRC1 expression in tumour epithelial cells, we next assessed whether this transcriptional axis holds prognostic value in PCa patients.

Stratification of patients based on combined PGC1α and CTHRC1 expression revealed that PCa tumours with high PGC1α and low CTHRC1 mRNA levels exhibited significantly improved clinical outcomes compared to those with low PGC1α and high CTHRC1 expression. This association was consistently observed across two publicly available PCa cohorts and was further validated in a third independent patient cohort (Figure 3 F).

### CTHRC1 promotes cell–ECM adhesion, ECM organization, and tumour growth in prostate cancer

Having identified CTHRC1 as a candidate downstream effector of PGC1α with clinical relevance, we next investigated its functional role in PCa progression and ECM regulation. CTHRC1 is a secreted glycoprotein whose expression is markedly increased across multiple tumour types, including PCa (26–28) and Supplementary Figure 4 A), although its mechanistic contribution to tumour aggressiveness remains unknown. In line with previous studies, silencing of CTHRC1 in two independent PCa cell lines (Figure 4 A–B) resulted in a reduction of cell proliferation and migration (Supplementary Figure 4 B–D). Notably our extended analysis showed that CTHRC1 depletion markedly impaired the ability of PCa cells to adhere to ECM substrates under both 2D (Figure 4 C) and 3D conditions (Figure 4 D), with a more pronounced effect in DU145 cells than in PC3 (Figure 4 C-D And Supplementary Figure E). Importantly, the oncogenic phenotype associated to high CTHRC1 levels was confirmed *in ovo*, as CTHRC1 silencing led to a marked reduction in tumour formation (Figure 4 E). Moreover, analysis of ECM organization revealed that tumours with low CTHRC1 expression were characterized by disorganized and isotropic collagen fibres, whereas CTHRC1-high tumours displayed highly aligned collagen structures (Figure 4 F), indicative of a more aggressive phenotype.

**Figure 4.**
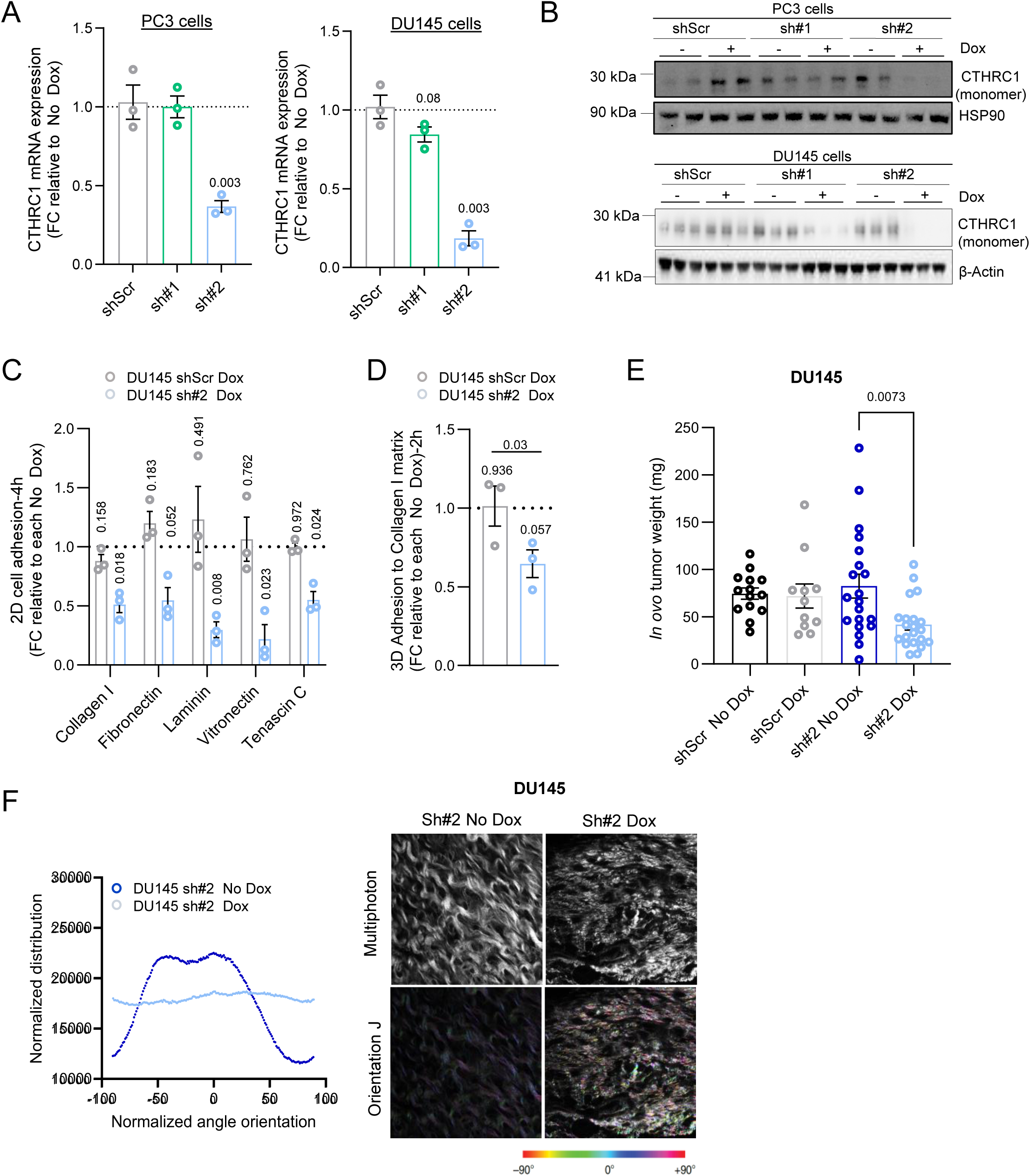
CTHRC1 silencing reduce prostate cancer cell–ECM interactions and ECM organisation. **A-B.** Validation of CTHRC1 silencing using two independent short hairpin RNAs and a scramble control in PC3 and DU145 cells by RT-qPCR (A, n=3) and by Western blot (B, n=3). **C**. Effect of CTHRC1 silencing in the capacity of DU145 cells to adhere to a panel of 2D extracellular matrixes *in vitro* (n=3). **D**. Effect of CTHRC1 silencing in the capacity of DU145 cells to adhere to 3D collagen-rich ECM in vitro (n=3). **E**. Effect of CTHRC1 silencing in the capacity of DU145 to form tumours *in ovo* (n= 21-14 tumours). **F**. Quantification of collagen fibre alignment (left panel) in tumours generated *in ovo* by CTHRC1-expressing and non-expressing DU145 cells. Right panel correspond to representative second harmonic generation two-photon microscopy images of the aforementioned *in ovo* tumours. Scale bar represents 20µm and colour coding represents angle degree. Dox: doxycycline; FC: fold change. Statistics: One sample t-test with reference value 1 (A, C and D) and unpaired t-test (E). Numbers represent p-values. Error bars indicate s.e.m.

Taken together, these data demonstrate that CTHRC1 promotes cell–ECM adhesion and ECM organization and phenocopies the effects of PGC1α loss, supporting its role as a key downstream effector of the PGC1α-regulated ECM program.

### CTHRC1 is necessary and sufficient to mediate PGC1α-dependent adhesion and extracellular matrix remodelling

We next sought to determine whether CTHRC1 is functionally required and sufficient to mediate the PGC1α-dependent ECM altered phenotype. To this end, we restored CTHRC1 expression in PC3 cells using a doxycycline-inducible system enabling co-expression with PGC1α (Supplementary Figure 5 A). Rescue efficiency was validated both intracellularly (Supplementary Figure 5 B–C) and at the level of secretion (Supplementary Figure 5 D-E).

Re-expression of CTHRC1 fully rescued the impaired adhesive capacity of PGC1α-expressing cells across multiple ECM substrates under both 2D (Figure 5 A) and 3D conditions (Figure 5 B), demonstrating that CTHRC1 is sufficient to restore cell–ECM interactions.

**Figure 5.**
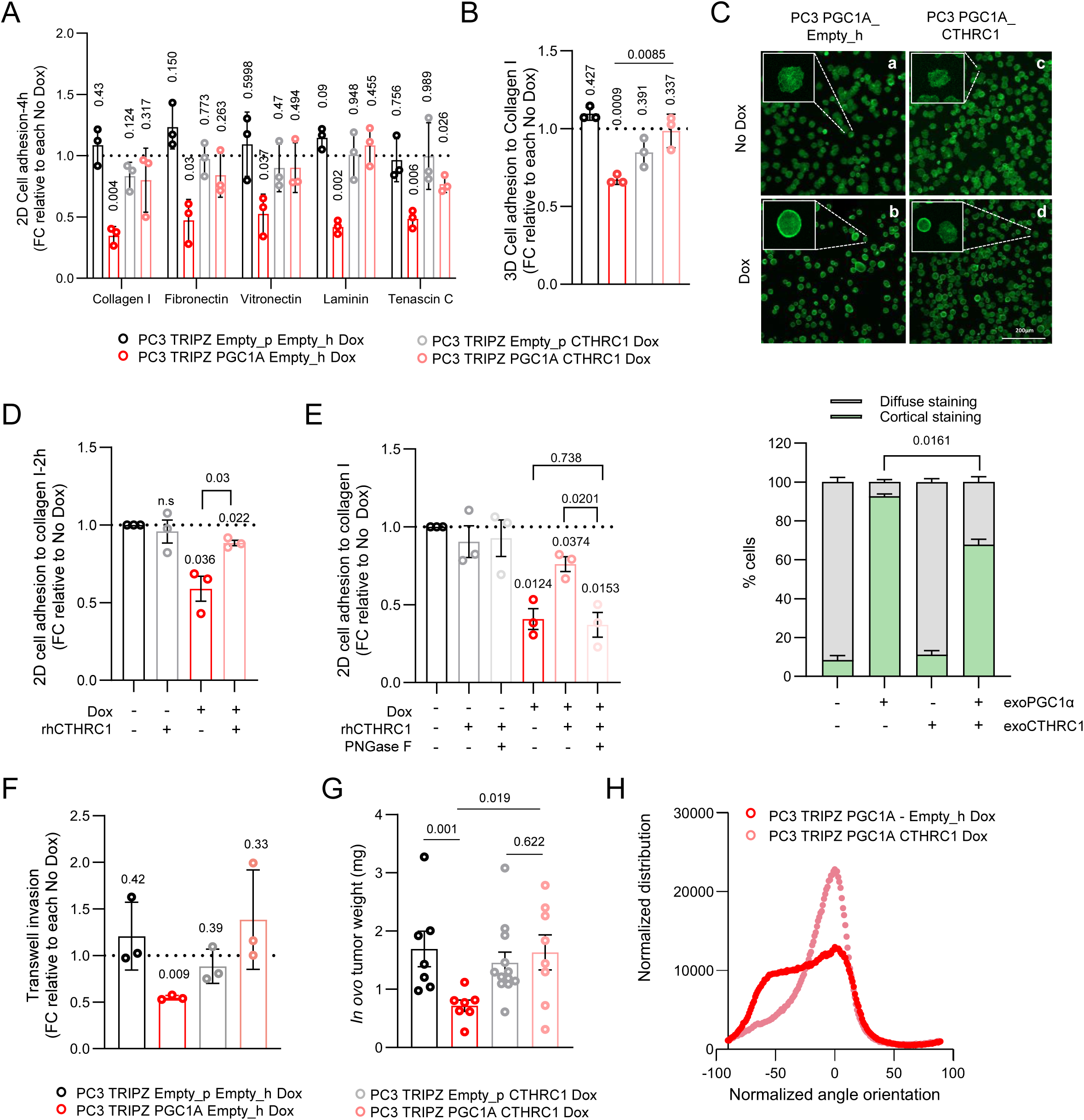
CTHRC1 is necessary and sufficient to mediate PGC1α-dependent phenotype. **A-B.** Effect of CTHRC1 rescue in the capacity of PGC1α-expressing and non-expressing PC3 cells to adhere to a panel of 2D extracellular matrixes (A) and 3D collagen-rich ECM (B) *in vitro* (n=3). **C**. Phalloidin staining showing differential F-actin distribution in PGC1α-expressing and non-expressing PC3 cells with or without rescued expression of CTHRC1. Right panel represents the % of cells with cortical and diffuse F-actin distribution in each cell line (n=3). **D**. Effect on 2D cell adhesion to collagen I of PGC1α-expressing and non-expressing cells treated with recombinant CTHRC1 protein (28 pg/mL) and control. **E**. Effect on 2D cell adhesion to collagen I of PGC1α-expressing and non-expressing cells treated with CTHRC1 recombinant protein and glycosylation inhibitor. **F.** Transwell invasion experiments in PGC1α-expressing and non-expressing with and without the re-expression of CTHRC1. **G.** *In ovo* tumour formation of PGC1α-expressing and non-expressing with and without the re-expression of CTHRC1. **H**. Multiphoton imaging analysis of the previously mentioned in ovo tumours. Dox: doxycycline; FC: fold change; Empty_p= control plasmid for puromycin selection; Empty_h= control plasmid for hygromycin selection. Statistics: One sample t-test with reference value 1 (A, B, F and E), paired t-test (B, C, D and E only represented the comparison between PGC1α +Dox vs PGC1α CTHRC1 + Dox conditions) and unpaired t-test (G). Numbers represent p-values. Error bars indicate s.e.m.

In agreement with these changes, CTHRC1 appears to contribute to the regulation of early cytoskeletal dynamics during cell adhesion. In its reduced state (PGC1α-expressing cells), PCa cells exhibited cortical F-actin redistribution forming ring-like structures upon adhesion to collagen I, whereas control cells displayed a diffuse phalloidin staining pattern consistent with active actin remodelling (Figure 5 C, panels a–b). Notably, CTHRC1 re-expression reverted this phenotype, restoring a diffuse actin organization comparable to control cells (Figure 5 C, panels c–d), thereby linking CTHRC1 downregulation to impaired cytoskeletal remodelling.

To determine whether the secreted form of CTHRC1 was sufficient to mediate the PGC1α-dependent adhesion phenotype, we complemented the genetic approach with functional rescue experiments using CTHRC1-recombinant protein. Based on ELISA quantification of secreted CTHRC1 (Supplementary Figure 5 D), we defined 28 pg/mL as the concentration required to restore basal extracellular levels in PC3 cells. Treatment of PGC1α-expressing cells with recombinant CTHRC1 was sufficient to rescue adhesion to collagen I in 2D assays (Figure 5 D), recapitulating the effects observed upon genetic restoration. These findings indicate that the impaired adhesive capacity induced by PGC1α is mediated, at least in part, through reduced levels of secreted CTHRC1.

Given that CTHRC1 is a glycosylated protein and that glycosylation is known to regulate the function and organization of ECM components (29), we next assessed the contribution of this modification. Deglycosylation of recombinant CTHRC1, confirmed by western blot (Supplementary Figure 5 F), completely abolished its ability to rescue the adhesion defect in PGC1α-expressing cells (Figure 5 E), demonstrating that glycosylation is required for the pro-adhesive function of CTHRC1. These findings provide mechanistic insight into how CTHRC1 regulates cell–ECM interactions at the molecular level.

Notably, co-expression of CTHRC1 in PGC1α-expressing cells significantly increased their invasive capacity (Figure 5 F) and restored tumour growth *in ovo* compared to PGC1α expression alone (Figure 5 G). Moreover, collagen fibres in these tumours reverted from a random aligned phenotype observed in PGC1α-expressing tumours to an anisotropic architecture of the fibres upon CTHRC1 restoration (Figure 5 H). Collectively, these data showed that downregulation of CTHRC1 is necessary and sufficient to mediate PGC1α-dependent alterations in cell adhesion and ECM organization, with its secreted and glycosylated form playing a central mechanistic role.

Overall, our results demonstrate that loss of PGC1α drives early ECM remodelling in PCa, characterized by transcriptional and structural alterations that promote tumour aggressiveness. Through integrated *in vivo* and multi-omics approaches, we identified CTHRC1 as a clinically relevant key PGC1α-regulated matrisome effector that controls cell–ECM interactions, collagen organization, and tumour growth. Gain-and loss-of-functional studies establish CTHRC1 as both necessary and sufficient to mediate PGC1α-dependent ECM phenotypes. Importantly, this activity depends on its secreted, glycosylated form, underscoring a critical post-translational requirement for its function.

## DISCUSSION

Tumour progression is increasingly recognized as a process shaped not only by cell-intrinsic alterations but also by dynamic interactions between cancer cells and the extracellular matrix (ECM) (30, 31). Here, we identify loss of PGC1α as an early epithelial event that drives ECM remodelling in prostate cancer (PCa), thereby extending its tumour-suppressive role beyond metabolic regulation (6, 9, 32–36). Our data demonstrate that PGC1α loss activates transcriptional programs that promote collagen deposition, alter ECM architecture, and enhance tumour aggressiveness prior to metastatic dissemination, positioning PGC1α as a central regulator linking epithelial transcriptional states with microenvironmental remodelling.

Mechanistically, our findings place ECM regulation among the dominant processes repressed by the PGC1α–ERRα axis. While this axis has been primarily associated with metabolic control (37), our results reveal a broader function in constraining tumour progression through transcriptional repression of ECM-related programs. In this context, ERRα emerges as a key mediator of this repressive activity (38), coordinating gene networks involved in cell–matrix interactions and extracellular signalling. These data support a model in which PGC1α acts as a transcriptional hub integrating intracellular metabolic cues with extracellular structural regulation, further expanding the emerging paradigm that metabolic regulators can directly influence extracellular matrix biology (39, 40).

A major advance of this study is the identification of CTHRC1 as a critical downstream effector of PGC1α. We show that CTHRC1 expression is tightly repressed by PGC1α in tumour epithelial cells and that epithelial-derived CTHRC1 contributes functionally to ECM remodelling, complementing its previously described roles in cancer, fibrosis and stromal biology (28, 41–44). Functional analyses demonstrate that CTHRC1 is both necessary and sufficient to restore ECM adhesion, cytoskeletal dynamics, collagen organization, and tumorigenic capacity downstream of PGC1α. Notably, we uncover that the extracellular, glycosylated form of CTHRC1 is required for its pro-adhesive function, highlighting a mechanistic role for post-translational modifications in ECM-associated signalling (15, 45).

Importantly, our data establish CTHRC1 as a causal mediator of the ECM alterations observed in PGC1α-deficient tumours. Restoration of CTHRC1 is sufficient to rescue adhesion signalling and re-establish aligned collagen architectures associated with aggressive disease. These data support existing literature (46–48) and demonstrate that modulation of a single ECM-associated factor can reprogram both tumour cell behaviour and microenvironmental structure in vivo.

Beyond providing mechanistic insights, our study has important clinical implications as ECM remodelling and collagen alignment emerge early during prostate tumorigenesis following PGC1α loss and are recapitulated in human tumours. These structural features have been associated with poor prognosis across multiple cancer types (49–53). Notably, the combined expression of PGC1α and CTHRC1 improves patient stratification across independent cohorts, supporting the prognostic value of this axis. While further validation—particularly at the protein level—is required, these findings highlight the PGC1α–CTHRC1 axis as a mechanistic and clinically relevant regulator of PCa aggressiveness.

In summary, our study uncovers an epithelial transcriptional program linking PGC1α loss to ECM remodelling and identifies the PGC1α–CTHRC1 axis as a key driver of tumour progression. These findings provide new insight into early events shaping tumour aggressiveness and highlight transcriptional control of the matrisome as a critical determinant of PCa biology.

## MATERIALS AND METHODS

### Reagents

Doxycycline hyclate (Sigma #D9891) was applied to induce or suppress gene activity in tetracycline-regulated constructs. After lentiviral infection, cells were maintained under selection using Puromycin (Sigma #P8833) and Hygromycin B (Sigma #SBR00039).

### Animals

All mouse experiments were carried out following the ethical guidelines established by the Biosafety and Welfare Committee at CICbioGUNE. The procedures employed were carried out following the recommendations from Association for Assessment and Accreditation of Laboratory Animal Care International. GEMM experiments were generated and carried out as reported in a mixed background (8). The *Pten loxP* and *Pgc1a loxP* conditional knockout alleles have been described elsewhere (54). Prostate tumours were isolated from three-month *Pten^pc-/-^ Pgc1a^pc+/+^* and *Pten^pc-/-^ Pgc1a^pc-/-^* mice. Following the ethical guidelines, mice were sacrificed and the prostate (anterior, ventral and dorso-lateral lobes) was extracted and divided in two sections: (1) snap-frozen in liquid nitrogen and stored at −80 °C for further molecular analysis and (2) fixed in formalin for histopathological studies.

Orthotopic xenograft experiments were performed as described previously (55), injecting 1 x 10^6^ cells in the ventral prostate lobe of 6 weeks old Hsd:AthymicNude-Foxn1nu “nude” mouse (Envigo). Animals were assigned to chow or doxycycline diet regime (Research diets, D12100402) 1 day after the inoculation. PC3-luc growth was tracked by bioluminescence imaging using IVIS technology (PerkinElmer). Retro-orbital injections of 50 µL luciferin (15 mg/mL; PerkinElmer) were administered before imaging during the follow up and a volume equal to the weight of animal multiplied by 5 at end point (30 mg/mL; PerkinElmer).

### Patients

Disease free-survival, correlation and gene expression analysis were performed using PCa patient’s data obtained from publicly available databases from cBiorportal or Cancertool (56) websites or from a local PCa cohort previously described (23).

### Histopathological analysis

For paraffin embedded tissues were stained with Picro Sirius Red Stain Kit (#ab150681; collagen I and III) following manufacturer guidelines. Briefly, paraffin-embedded tissue sections were deparaffinised and hydrated in distilled water. Sections were then incubated with Picro Sirius Red solution for 60 minutes at room temperature. Slides were rinsed twice in 0.5% acetic acid solution, followed by rinsing in absolute ethanol. Sections were then dehydrated in two changes of absolute ethanol, cleared, and mounted using synthetic resin. Stained sections were imaged using light microscopy.

### Chorioallantoic membrane assay (CAM assay)

Chorioallantoic membrane (CAM) assays were performed as previously described (12). Briefly, fertilized chicken eggs (Santa Isabel farm, Córdoba, Spain) were inoculated at embryonic day 10 with 1 × 10⁶ cells resuspended in serum-free DMEM. Eggs were incubated for 5 days, and tumours were excised at embryonic day 15, weighed, and processed for downstream analyses.

### Tissue decellularisation

Decellularisation protocol of tumour samples was carried out using a multi-step decellularisation process. Briefly, freshly collected tissues were immersed in 1 mL of decellularisation buffer 1 [10 mM Tris-HCl (pH 8.0), 0.1% EDTA, and 10 KIU/mL aprotinin (Roche #10236624001)] and incubated under continuous agitation at 4 °C for 48 h. Subsequently, the medium was replaced with decellularisation buffer 2 [same composition of buffer 1 supplemented with 3% Triton™ X-100] and maintained under agitation at 4 °C for an additional 72 h. After this period, samples were transferred to 1.5 mL tubes and washed with PBS supplemented with 2% penicillin–streptomycin. To ensure complete removal of genetic material, tissues were incubated at 37 °C for 24 h in a nuclease solution (PBS with 2% penicillin–streptomycin) containing RNase at 2 µg/mL and DNase at 200 µg/mL.

### Multiphoton microscopy

Multiphoton imaging was performed using a Zeiss LSM 880 NLO multiphoton confocal microscope equipped with a Mai Tai DeepSee tunable femtosecond laser (690–1040 nm), located at the CIC biomaGUNE microscopy facility. Samples were imaged using a 20× objective lens (NA 0.8). Multiphoton excitation was set at 820 nm. Signal from collagen fibres was collected using non-descanned detectors (NDD, Big2 module) in the 380–430 nm range (channel 1), while unspecific tissue autofluorescence was simultaneously detected in the 465–515 nm range (channel 2). Images were acquired at a resolution of 1024 × 1024 pixels, with a pixel size of 0.69 µm. The pixel dwell time was set to 4 µs, and each image represents the average of two scans. All acquisitions were performed using Zeiss ZEN Black software.

Collagen fibre alignment was quantified using the OrientationJ plugin (Biomedical Image Group, EPFL, Switzerland) implemented in ImageJ/Fiji. OrientationJ Analysis was performed using a local window (σ) of 2 pixel and a cubic spline gradient to generate color-coded orientation maps. OrientationJ Distribution was subsequently used to obtain orientation histograms and alignment statistics, which were normalized to the maximum angle (12).

### Immunophenotyping and Flow cytometry

Prostate tumours were harvested from 3-months old *Pten^pc-/-^ Pgc1a^pc+/+^* and *Pten^pc-/-^Pgc1a^pc-/-^* mice and digested with 1 mg/mL collagenase (37°C 1hr at 900 rpm). The digested suspension was centrifuged for 6 min at 350 x g and 4°C. The pellet was re-suspended in PBS and passed through nylon filters to remove clamps. The resulting single-cell suspension was plated in V-bottom 96-well plates (ThermoFisher, Cat. #: 249570). Samples were first incubated with a fixable LIVE/DEAD-BB515 viability dye (ThermoFisher, Cat. #: L34960) for 30 min at 4 °C, protected from light. Fc receptors were subsequently blocked using TruStain FcX (1:50, BioLegend, Cat. #: 101320) for 10 min at room temperature in the dark. Cells were then stained with fluorochrome-conjugated antibodies diluted in staining buffer (ThermoFisher, Cat. #: 00-4222-26) for 30 min at 4 °C, protected from light. To determine absolute cell numbers, CountBright™ Absolute Counting Beads (ThermoFisher, Cat. #: C36995) were added to each sample prior to acquisition, and cell counts were calculated based on the ratio between bead events and cellular events. Acquisition was performed using BD FACSymphony flow cytometer and data was analysed using FlowJo v10 software.

The antibodies used were: CD3 – BUV 737 (BD Biosciences, 612803), CD4 – BUV 395 (BD Biosciences, 563790), CD8 – BUV 563 (BD Biosciences, 748535), CD11b – BUV805 (BD Biosciences, 741934), CD11c – PE (Biolegend, 117307), CD19 – APC (Biolegend, 115512), CD45.2 – BV480 (BD Biosciences, 566073), F4/80 – APC Fire 750 (Biolegend, 123151), Gr1 – BV711 (Biolegend, 108443), NK 1.1 – BV 605 (Biolegend, 108753).

### Cell culture

Human prostate carcinoma cell lines PC3 (RRID:CVCL_0035) and DU145 (RRID:CVCL_0105) were obtained from the Leibniz Institute DSMZ (Deutsche Sammlung von Mikroorganismen und Zellkulturen GmbH), which provided certificates of authentication. Cell line identity was periodically confirmed by microsatellite-based profiling, and none of the cell lines used in this study were listed in databases of commonly misidentified cell lines curated by the International Cell Line Authentication Committee and NCBI Biosample. All cultures were routinely tested and confirmed negative for Mycoplasma contamination. Both cell lines were maintained in Dulbecco’s Modified Eagle Medium (DMEM) without pyruvate, supplemented with 10% (v/v) fetal bovine serum (FBS) and 1% (v/v) penicillin–streptomycin. Cells were cultured at 37 °C in a humidified incubator with 5% CO₂.

None of the cell lines used in this study were found in the database of commonly misidentified cell lines maintained by the International Cell Line Authentication Committee and NCBI Biosample. All cell lines were routinely monitored for Mycoplasma contamination. Cells were transduced with a modified TRIPZ (Dharmacon) doxycycline inducible lentiviral construct conferring puromycin resistance, in which the RFP and miR30 region were removed and replaced by HA-Flag-Pgc1a or HA-Flag-CTHRC1. Lentiviral particles were generated in HEK293FT (RRID:CVCL_6911) cells using standard protocols. Viral supernatants obtained after packaging were used to infect target cells. To rescue CTHRC1 expression, PGC1a-expressing cells were transduced with TRIPZ-Ha-Flag-CTHRC1 vector in which the puromycin resistance cassette was replaced with the hygromycin one. In parallel, empty vector PC3 cells were also generated as controls.

For ESRRA deletion, single-guide RNA (sgRNA) constructs targeting ESRRA were used as previously described (9).

For gene expression silencing of CTHRC1, DU145 and PC3 cells were transduced with the doxycycline-inducible lentiviral plkoTet-ON vectors (modified from Addgene #21915) encoding 2 independent shRNA (TRCN0000062245 and TRCN0000371999 Sigma-Aldrich) targeting CTHRC1 mRNA and with the scramble control. Cells were then selected with puromycin (2 μg/mL) for 3 days.

### Cellular assays

For doxycycline-inducible experiments, cells were pre-induced with doxycycline (0.5 µg/mL) for 3 days prior to seed the experiments. A second doxycycline induction was performed when required.

Cell proliferation was quantified by crystal violet staining as previously described (8). Briefly, plates were fixed at the indicated time points with 10% formalin, washed with 1X PBS, and stained with crystal violet solution (0.1% crystal violet in 20% methanol) for 1 h. Plates were air-dried, scanned, and crystal violet precipitates were solubilized in 10% acetic acid for 30 min. Absorbance was measured at 600 nm using a GloMax® Discover Microplate Reader (GM3000).

For adhesion assays, 60,000 cells per well were seeded in 24-well plates pre-coated with the indicated extracellular matrix proteins: rat tail collagen I (Corning #354236, 50 µg/mL diluted in 0.02 N acetic acid), fibronectin (5 µg/mL in PBS Sigma #F2006), laminin (3 µg/mL in PBS Sigma #L6274), vitronectin (5 µg/mL in PBS Sigma #SRP3186), or tenascin C (0,5 µg/mL in mqH_2_0 Sigma #CC065), independently. After, plates were washed with PBS, fixed with 10% formalin, and stained with crystal violet as described above.

Three-dimensional collagen I matrices were generated using type I bovine collagen solution (Advanced BioMatrix PureCol® #5005). Collagen matrices were prepared according to the manufacturer’s instructions (neutralized with NaOH to physiological pH). Cells were seeded onto the matrices and then fixed with 10% formalin as described above.

Transwell invasion assays were performed using Matrigel-coated chambers (BD BioCoat, #354480). 50,000 cells per well were resuspended in DMEM without FBS and seeded into the upper chamber. A total of 1.4 mL of complete DMEM was added to the lower chamber. Cells were incubated at 37 °C and 5% CO₂ for 48 hours. Invasion was stopped by washing the inserts twice with 1X PBS and removing non-invading cells from the upper surface of the membrane using a cotton swab. Membranes were fixed with 10% formalin for 15 minutes at 4 °C and stained with crystal violet.

Migration assays were performed using migration chambers (Corning, #351185) containing polycarbonate membranes with 8 μm pores. Cell seeding, washing, and fixation conditions were the same as those used for the invasion assay, but the incubation time was 24 hours. Images were acquired using a light microscope, and migrated cells were quantified using Fiji software following the same procedure used for invasion assays.

Bromodeoxyuridine (BrdU; 5′-bromo-2′-deoxyuridine) incorporation. 15,000 DU145 cells were seeded onto coverslips placed in 24-well plates. Four days after seeding, BrdU (Sigma-Aldrich, B9285) was added dropwise to each well at a final concentration of 3 µg/µl, and cells were incubated for 4 h. The culture medium was then removed, cells were gently washed with PBS and fixed with 4% PFA in PBS for 15 min at room temperature. After fixation, coverslips were washed with PBS and stored at 4 °C until further processing. For BrdU immunostaining, DNA denaturation was performed by incubating coverslips with 2 M HCl for 5 min, followed by neutralization with 0.1 M sodium borate for 5 min and subsequent PBS washes. Cells were permeabilised with 0.1% Triton X-100 and glycine in PBS for 5 min and then blocked with 10% goat serum in PBS (Vector Laboratories, S-1000-20) for 30 min at room temperature. Coverslips were incubated overnight at 4 °C in a humidified chamber with an anti-BrdU primary antibody (BD Biosciences, 555627) diluted 1:100 in blocking solution. The following day, after washing with PBS, coverslips were incubated with the appropriate Alexa Fluor–conjugated secondary antibody (mouse Alexa Fluor 488 or 568; Thermo Fisher Scientific, A-21121 or A-21144) diluted 1:1000 in blocking solution for 2 h at room temperature in the dark. After PBS washes, coverslips were mounted using ProLong™ Gold Antifade Mountant with DAPI (Thermo Fisher Scientific, P36931). Fluorescence images were acquired using a fluorescence microscope, and BrdU-positive cells were quantified using ImageJ software.

For visualisation of F-actin fibres, fixed cells were stained with phalloidin (Invitrogen #A12379) and nuclei were counterstained using DAPI mounting medium. Images were acquired using a Zeiss LSM800 confocal microscope.

Secreted CTHRC1 levels were quantified using a human CTHRC1 ELISA kit (Abcam, #ab274399) according to the manufacturer’s instructions. Standards and samples were added to a pre-coated 96-well plate and incubated following the manufacturer’s protocol. After washing, detection antibody and HRP-conjugate were added, and signal was developed using TMB substrate. Absorbance was measured at 450 nm with wavelength correction at 570 nm.

Recombinant human CTHRC1 protein (MedChemExpress, #HY-P70092) was reconstituted and handled according to the manufacturer’s instructions.

Deglycosylation of 1 μg of recombinant CTHRC1 protein was performed using PNGase F (New England Biolabs, #P0704; 0.1 units) according to the manufacturer’s instructions.

### Molecular assays

For protein extraction, cells were seeded on 6-well plates and 3 days post-seeding, lysed using RIPA buffer (#sc-24948). Protein concentration was determined using the Pierce™ BCA Protein Assay Kit (Thermo Fisher Scientific #NP0007). Samples were prepared in 1X Läemmli buffer (from 5X stock: 10% SDS, 50 mM Tris pH 6.8, 50% glycerol, 1% β-mercaptoethanol, 0.01 M DTT, and 0.2 mg/mL bromophenol blue) and denatured at 95 °C for 5 min. Proteins were resolved using NuPAGE® Novex® 4-12% Bis-Tris gels in MOPS SDS buffer or Mini-Protean TGX Precast Gels in Tris-Glycine SDS buffer at 200 V. Gels were transferred to nitrocellulose membranes (Amersham Protran) at 80 V for 1hour. After blocking with 5% non-fat milk in TBS-T (0.01% Tween-20) for 1 hour, membranes were incubated with primary antibodies (1:1,000) against PGC1α (#sc-13067), ERRα (#cs-13826), GAPDH (#cs-2118), CTHRC1 (#ab85739), HSP90 (#cs4874) and β-Actin (Sigma #A5441). Mouse and rabbit secondary antibodies were purchased from Jackson ImmunoResearch. After standard SDS-PAGE and western blotting techniques, proteins were visualized using the ECL system in the iBright FL1000 Imaging System and BioRad.

Total RNA was isolated using the NucleoSpin® RNA isolation kit (Macherey-Nagel #740955.240C). Animal tissues were homogenized using a Precellys® Evolution Touch (Bertin technologies, #P002511-PEVT0-A.0) prior to RNA extraction. Total RNA was subsequently isolated using a trizol-based implementation of the NucleoSpin® RNA isolation kit protocol as previously reported (NCB, 2016). Complementary DNA (cDNA) was synthesized from 1 µg of total RNA using the Maxima H Minus cDNA Synthesis Kit with dsDNase (Thermo Scientific #M1682). Quantitative PCR was performed on a QuantStudio 5 (QS5) Real-Time PCR System (Applied Biosystems). Target gene expression was detected using PrimeTime™ TaqMan probes (Integrated DNA Technologies). Probe references: CTHRC1: Hs.PT.58.39259295; PPARGC1A: Hs.PT.58.14965839 Mm.PT.58.9404608; GAPDH: Hs.PT.39a.22214836; PLAU: Hs.PT.58.22549485; COL13A1: Hs.PT.58212348; IL1B: Hs.PT.58.151818.

### RNA sequencing and analysis

Quality of the tumour and culture cell extracted RNA samples was assessed using Qubit RNA HS Assay Kit and RNA integrity numbers (RINs) were obtained from Agilent 2100 Bioanalyze. Strand-specific RNA-seq libraries were generated using the Illumina TruSeq Stranded Total RNA Library Prep Kit according to the manufacturer’s instructions. High-throughput sequencing was performed on an Illumina NovaSeq 6000 platform, producing paired-end reads of 150 bp (2 × 150 bp) with an average depth of approximately 100 million reads per sample.

Raw sequencing reads were aligned to the reference genomes (hg38 and mm10, respectively) using STAR (version 2.7.5c) in two-pass mode following recommended best practices. Resulting BAM files were processed and managed using SAMtools (version 1.9) together with STAR utilities. Gene-level read counts were generated using featureCounts (version 2.0.3) from the Subread package based on the corresponding gene annotation.

Downstream analyses were performed in R (version 3.6.0). Differential gene expression analysis was conducted using the EdgeR and Limma packages. Count data were normalized using the trimmed mean of M values (TMM) method, and statistical testing for differential expression was performed using linear modelling approaches implemented in these packages. Genes with an adjusted *P* value (false discovery rate, FDR) below 0.05 were considered significantly differentially expressed.

Transcriptomic data has been deposited in GEO with accession number GSE329602.

CAF marker gene sets (REF) were mapped to DEGs using dplyr (CRAN package: 10.32614/CRAN.package.dplyr) and tidyr (CRAN package: 10.32614/CRAN.package.tidyr) for data processing. Heatmaps were generated using the ComplexHeatmap package (57, 58).

Functional enrichment analysis of the differential expressed genes was performed using g:GOst webtool available in g:Profiler (https://biit.cs.ut.ee/gprofiler/gost).

### Label-free proteomic analysis

Snap-frozen prostate tumours orthotopically generated from non-expressing and PGC1a-expressing PC3 cells were processes for proteomic analysis. Proteins were extracted using 7 M urea, 2 M thiourea, 4% CHAPS. Samples were incubated for 30 min at RT under agitation and digested following the filter-aided sample preparation (FASP) protocol described by Wisniewski and colleagues in 2009. Trypsin was added to a trypsin:protein ratio of 1:10, and the mixture was incubated overnight at 37 °C, dried out in a RVC2 25 speedvac concentrator (Christ), and resuspended in 0.1% formic acid (FA).

Processed samples were analysed in a timsTOF Pro with PASEF (Bruker Daltonics) coupled online to an Evosep ONE (Evosep) liquid chromatograph. The sample (200 ng) was directly loaded in a 15 cm Evosep endurance column (Evosep) and resolved using the 30 SPD protocol (approx. 44 min runs). Data was acquired using a Data-Indepent Acquisition (DIA) schedule.

DIA data was processed with DIA-NN software for protein identification and quantification using default parameters. Searches were carried out against a database consisting of Homo sapiens protein entries from Uniprot/Swissprot in library-free mode. Carbamidomethylation of cysteine was considered as fixed modification, and oxidation of methionine as variable modification. An FDR<1% at peptide level was applied as a significance cut-off. Data was loaded onto Perseus platform for data processing (log2 transformation, filtering, imputation) and statistical analysis (FDR-corrected Student’s t-test).

### Bioinformatic analysis and statistics

No statistical method was used to predetermine sample size. The experiments were not randomized. No inclusion/exclusion criteria were pre-stablished. The investigators were not blinded to allocation during experiments and outcome assessment. *n* values represent the number of independent experiments performed, the number of individual mice, or patient specimens. Survival curves were generated using Kaplan-Meier estimates and compared using log-rank tests. For each independent in vitro experiment, normal distribution was assumed, and one-sample *t*-test was applied for one-component comparisons with control and Student t test for two-component comparisons. A minimum of three independent experiments was performed. For in vivo experiments, a normality test was calculated and statistical test applied accordingly and a minimum of 5 animals per group was used. Two-tailed statistical analysis was applied for experimental design without predicted result, and one-tailed for validation or hypothesis-driven experiments. Outlier values were detected as values greater than +3 standard deviations from the mean, or less than -3 standard deviations. The confidence level used for all the statistical analyses was of 95% (alpha value ¼ 0.05). GraphPad Prism 10 software (RRID:SCR_002798) was used for statistical calculations.

## Acknowledgments

Apologies to those whose related publications were not cited because of space limitations. We gratefully acknowledge the Basque Biobank for Research (BIOEF) for the support with clinical samples and data management, the SGIker Analytical and High-Resolution Microscopy Core at EHU, the Microscopy and Advanced Bioimaging Core at Mount Sinai, the Torrano lab for valuable input, and the Bravo-Cordero lab members for their technical support. AG and AM were funded by FPI predoctoral fellowships from MICINN (PRE2018-083607 and PRE2022-101304); IH was funded by the Juan de la Cierva Incorporación program (IJC2019-040709-I); AC was supported by the Basque Department of Industry, Tourism and Trade (Elkartek), the MICINN (PID2022-141553OB-I0 (FEDER/EU); Fundación Cris Contra el Cáncer (PR_EX_2021-22), Severo Ochoa Excellence Accreditation (CEX2021-001136-S), European Training Networks Project (grant agreement no.: 955534), Fundación AECC (Excelencia 2024 call, Premetacan - EPAEC246710BIO), iDIFFER network of Excellence (RED2024-153635-T), and the European Research Council (Consolidator Grant 819242); JBC was supported by R01CA244780 (NIH/NCI), R61CA278402 (NIH/NCI), BC241026 (BCRP DOD), the Irma T. Hirschl Trust and the Emerging Leader Award from the Mark Foundation and the Tisch Cancer Institute National Institutes of Health (NIH) Cancer Center grant (P30-CA196521); LVJ was funded by the Juan de la Cierva Incorporación program (IJC2020-044958-I), Ramón y Cajal program RYC2023-042567-I and Fundación CRIS Contra el Cáncer PR_TPD_2022-04 in partnership with Fundación Adey. The work of V Torrano was supported by Ramón y Cajal Program RYC-2017-22295, national grants PID2021-123372OB-I00 and PID2024-155980OB-I00 from MICINN, Fundación Científica AECC LABAE211656TORR, the Basque Department of Industry, Tourism and Trade (ELKARTEK24/10), Consolidación Investigadora 2023 CNS2023-143848 and Fundación FERO. CIBERONC was cofunded with FEDER funds and funded by ISCIII.

## Author contributions

Conception and design: Torrano V

Development of methodology: Gonzalo-Paulano A, Macchia A, Valcarcel L, Gardoki J, Armendariz U, Lejona I, Egia-Mendikute L.

Acquisition of data (provided animals, acquired and managed patients, provided facilities, etc.): Loizaga A, Ugalde-Olano A, Carracedo A, Unzueta A, Mendizabal I, Azkalgorta M, Elortza F, Aransay AM, Palazón A, Bravo-Cordero J.

Analysis and interpretation of data (e.g., statistical analysis, biostatistics, computational analysis): Hermanova I, Rondon-Lorefice I, Mendizabal I, Azkalgorta M, Elortza F, Aransay AM,

Writing, review, and/or revision of the manuscript: Valcarcel L and Torrano V

Administrative, technical, or material support (i.e., reporting or organizing data, constructing databases): Lectez B, Martín-Martín N, Astobiza I.

Study supervision: Valcarcel L and Torrano V

