## Supplementary Figures for "Loss of PGC1α drives extracellular matrix remodelling in prostate cancer through CTHRC1"

*Supplementary Figure 1. Gonzalo et al.*

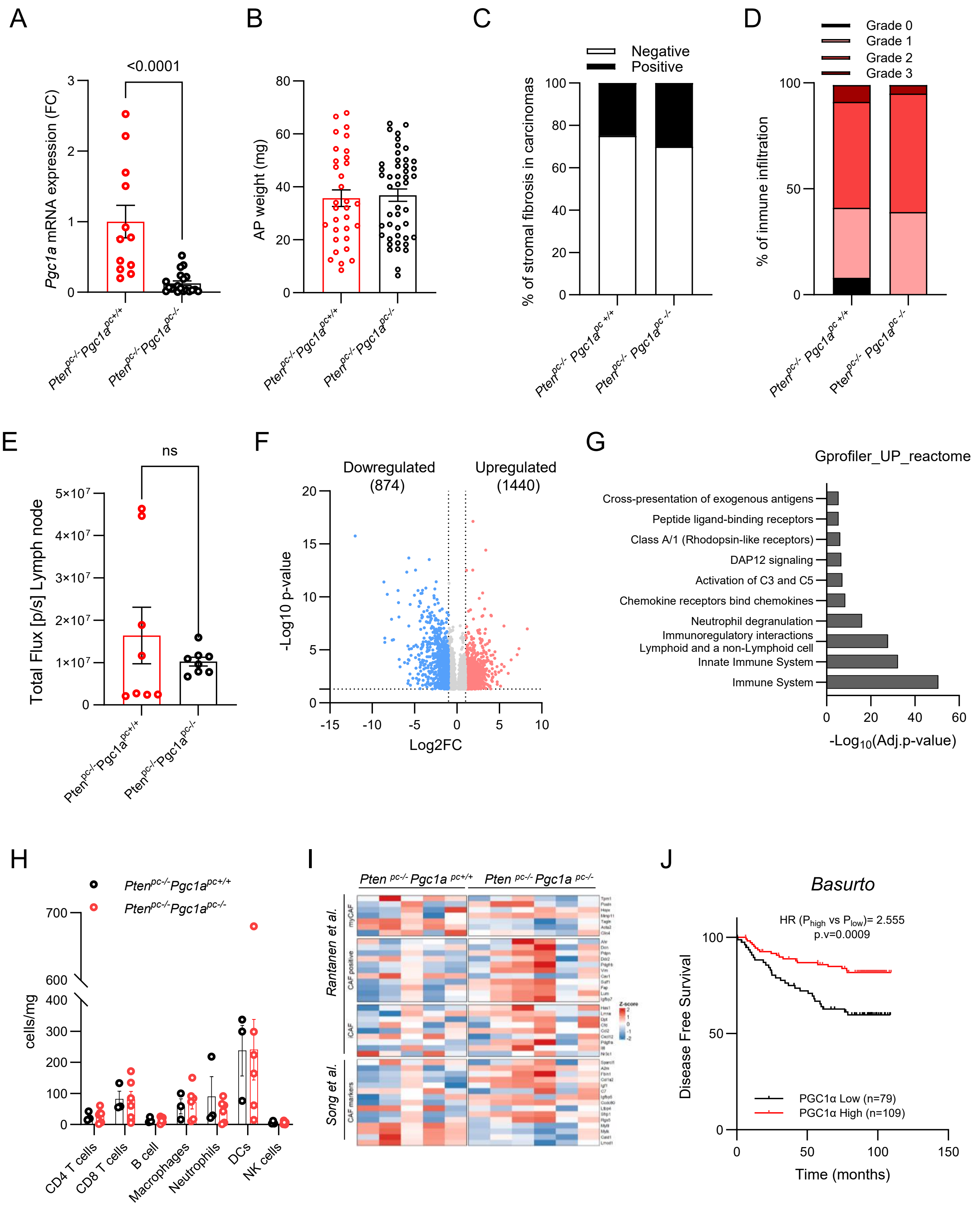

*Supplementary Figure 2. Gonzalo et al.*

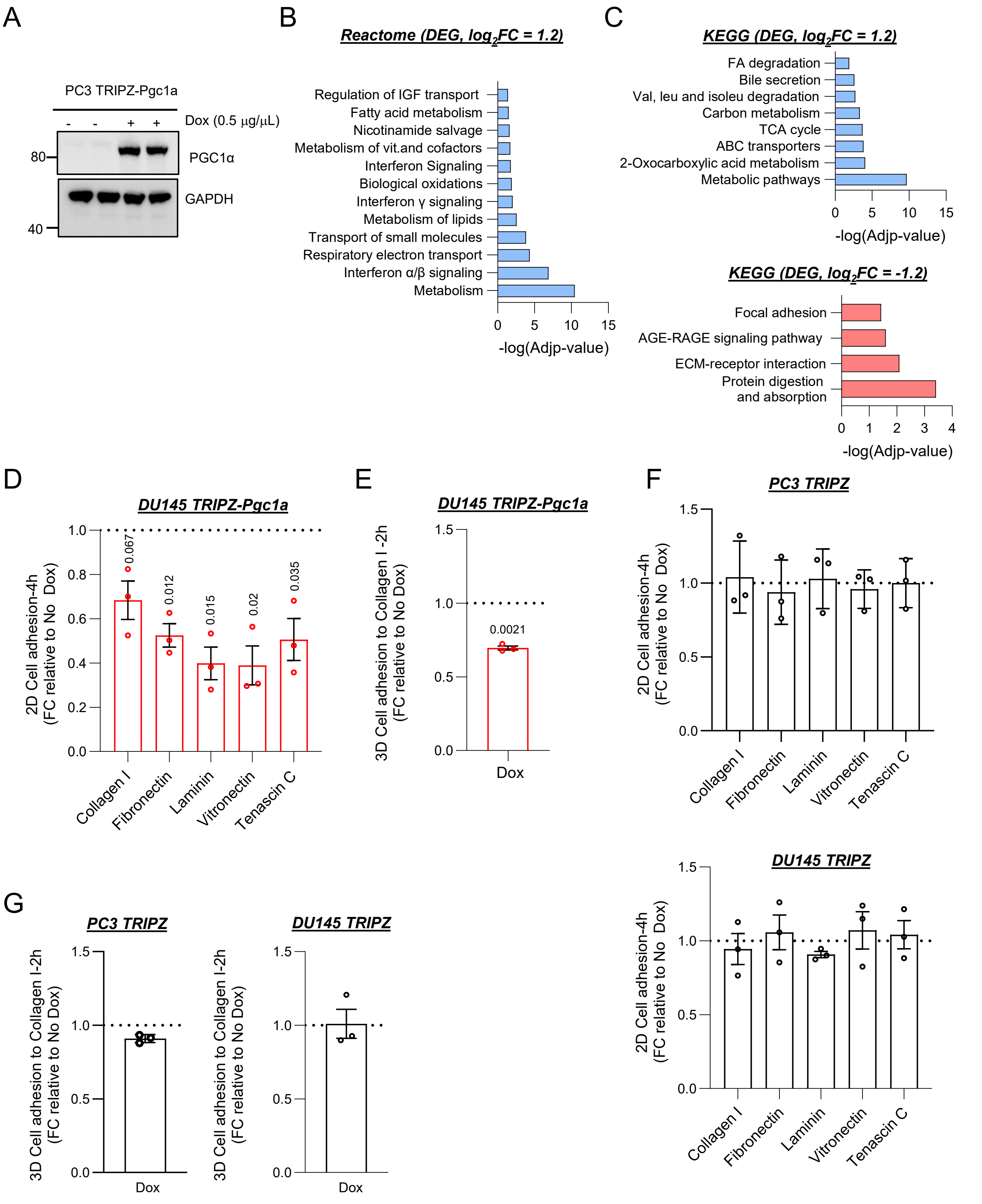

***Supplementary Figure 3. Gonzalo et al.***

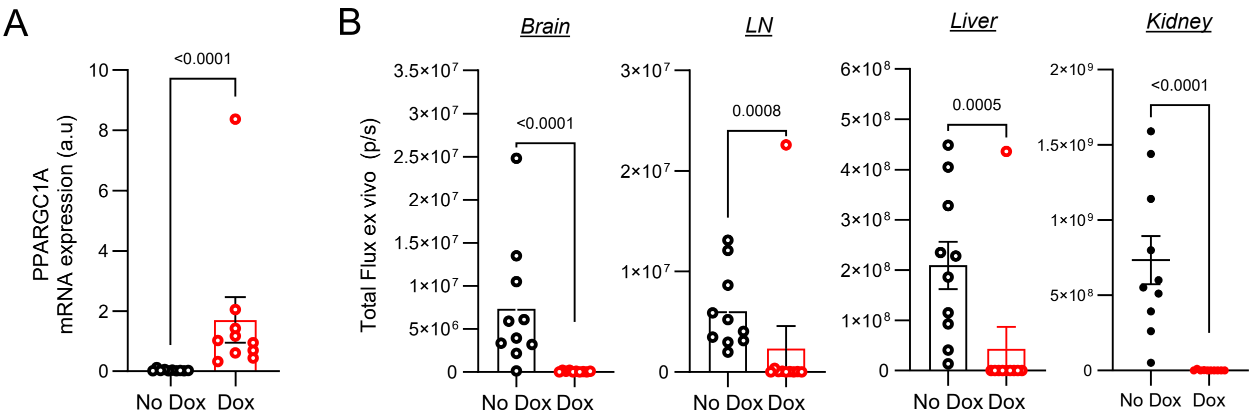


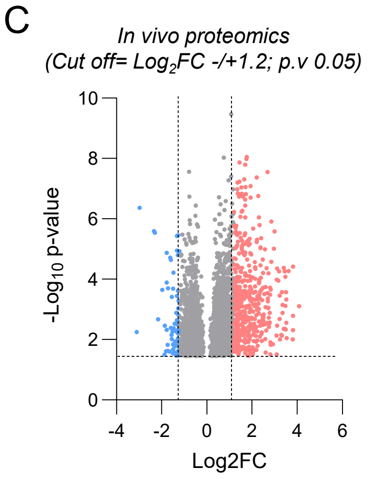


D

| | LC/MS | |
|---|---|---|
| | $Log_2FC$ | p-value |
| COL13A1 | -2.334 | 2.565E-06 |
| CTHRC1 | -1.498 | 2.063E-04 |
| IL1B | -1.777 | 1.345E-05 |
| PLAU | -1.623 | 1.931E-05 |
| | **RNAseq** | |
| | $Log_2FC$ | FDR |
| COL13A1 | -2.318 | 0 |
| CTHRC1 | -1.693 | 8.056E-215 |
| IL1B | -2.713 | 2.628E-97 |
| PLAU | -1.412 | 5.602E-157 |

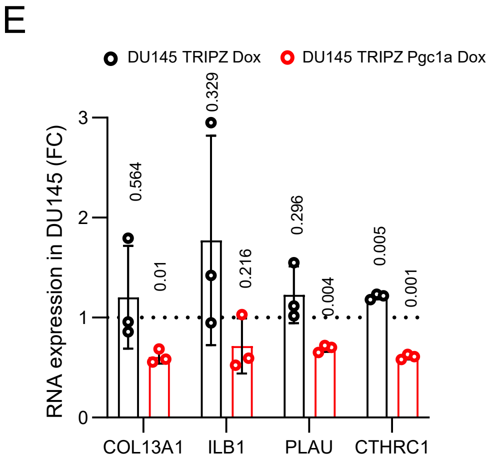


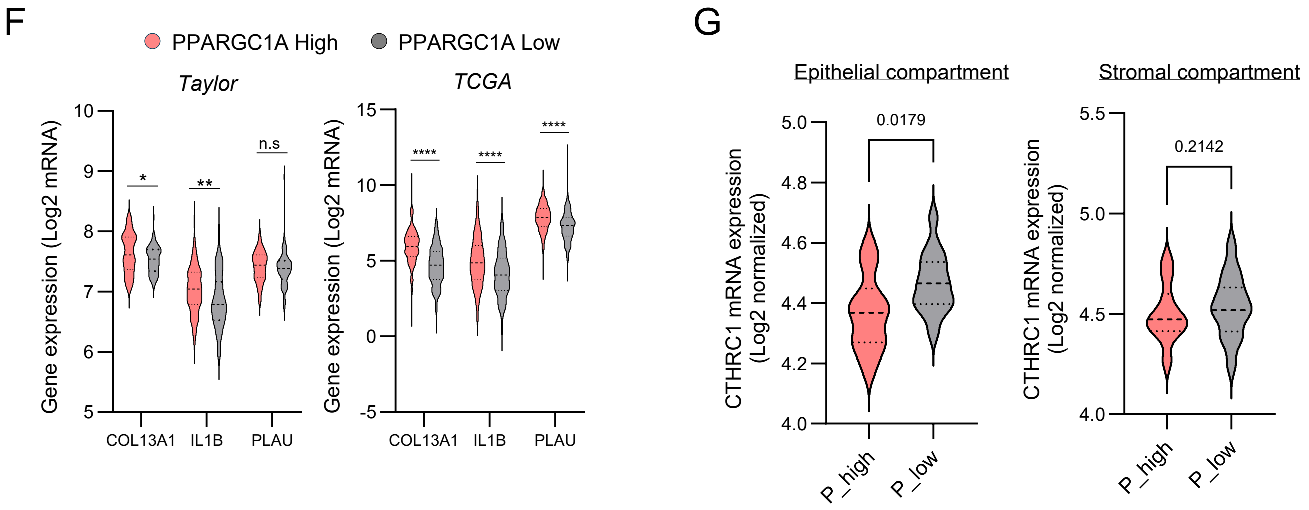

*Supplementary Figure 4. Gonzalo et al.*

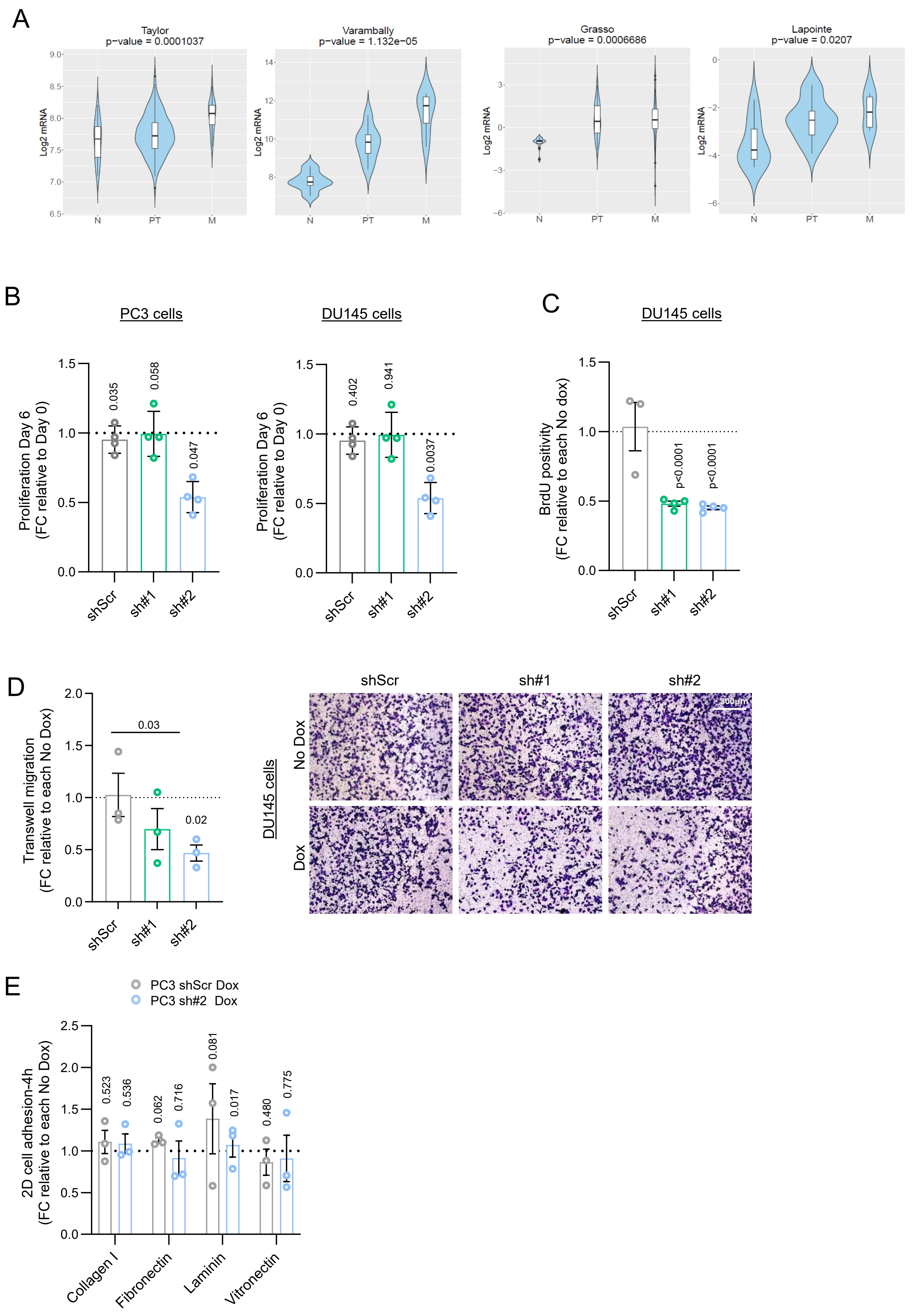

*Supplementary Figure 5. Gonzalo et al.*

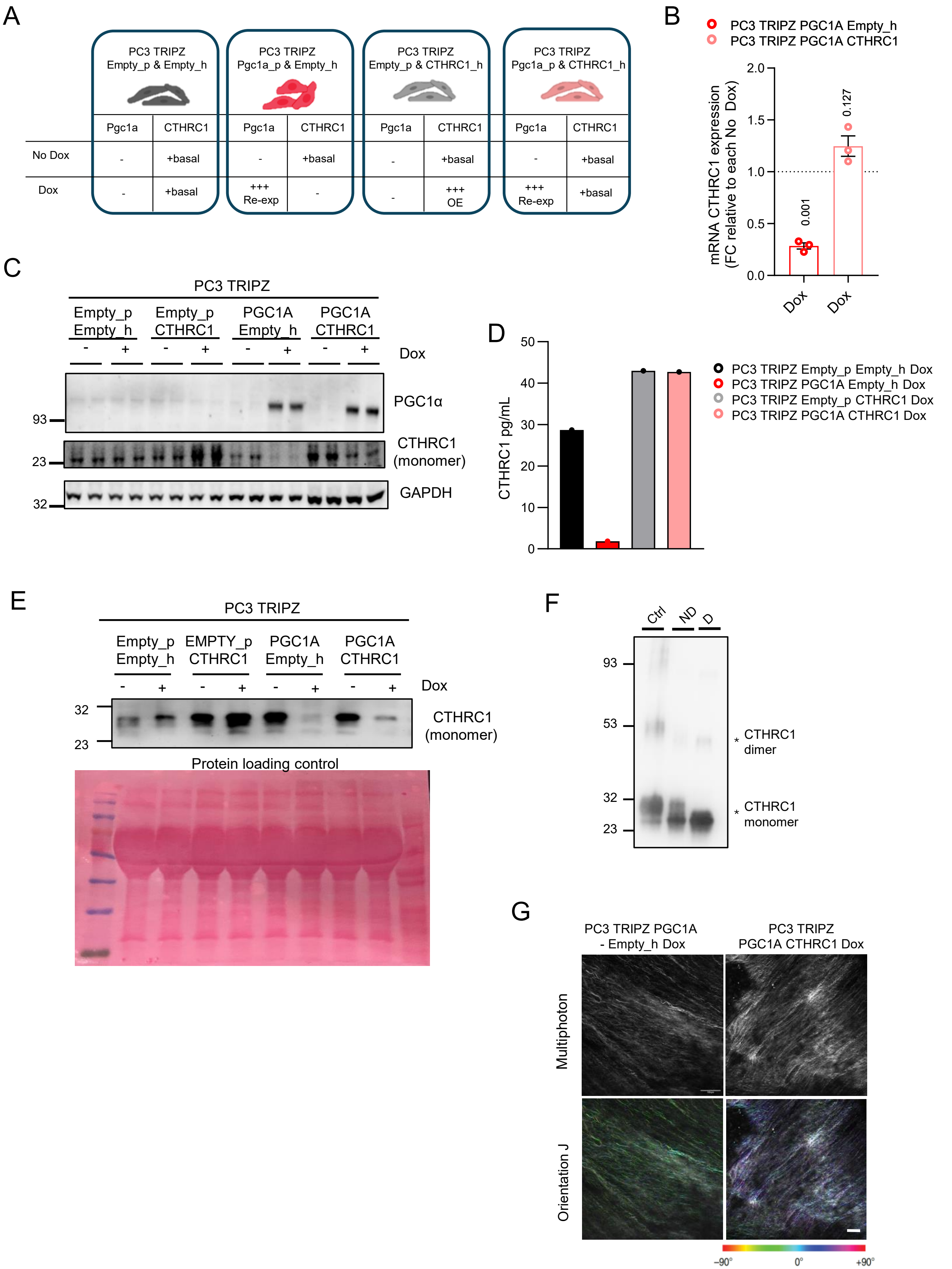
