## Supplementary Figure legends for "Loss of PGC1α drives extracellular matrix remodelling in prostate cancer through CTHRC1"

**Supplementary Figure 1. A.** Gene expression analysis (RT-qPCR) of Pgc1a in mouse prostate lobes (anterior prostate) (n=12). **B**. Anterior prostate weight in *Pten*$^{pc-/-}$ *Pgc1a*$^{pc+/+}$ and *Pten*$^{pc-/-}$ *Pgc1a*$^{pc-/-}$ mice, respectively. **C-D.** Stromal (E) and immune (F) infiltration quantifications based on hematoxylin-eosin staining of prostate tumours harvested from *Pten*$^{pc-/-}$ *Pgc1a*$^{pc+/+}$ and *Pten*$^{pc-/-}$ *Pgc1a*$^{pc-/-}$ 3-months old mice (n=12 and n=23, respectively). **E**. *Ex vivo* bioluminescence analysis of lymph nodes harvested from *Pten*$^{pc-/-}$ *Pgc1a*$^{pc+/+}$ Luc and *Pten*$^{pc-/-}$ *Pgc1a*$^{pc-/-}$-Luc 3-months old mice by means of total flux measurements (n=4 each genotype). **F.** Volcano plot representation of the differential gene expression analysis from *Pten*$^{pc-/-}$ *Pgc1a*$^{pc+/+}$ and *Pten*$^{pc-/-}$ *Pgc1a*$^{pc-/-}$ 3-months old mice transcriptomics (p-value < 0.05 and $Log_2$ FC -1/+1). **G.** Functional enrichment analysis (G-profiler-Reactome) of genes whose expression is upregulated upon Pgc1a deletion (and *Pten*$^{pc-/-}$ *Pgc1a*$^{pc-/-}$ versus *Pten*$^{pc-/-}$ *Pgc1a*$^{pc+/+}$ 3-months old mice). **H.** Differential quantification of immune cell populations in prostate tumours harvested from *Pten*$^{pc-/-}$ *Pgc1a*$^{pc+/+}$ (n=3) and *Pten*$^{pc-/-}$ *Pgc1a*$^{pc-/-}$ (n=6) 3-months old mice. Data is represented as the number of cells per mg of tissue. **I.** Heatmap representing the mRNA expression (data from RNAseq data-Supplementary table S1) of different cancer-associated fibroblast markers in *Pten*$^{pc-/-}$ *Pgc1a*$^{pc+/+}$ and *Pten*$^{pc-/-}$ *Pgc1a*$^{pc-/-}$ prostate tumours. **J.** Kaplan–Meier analysis of biochemical recurrence–free survival stratified by PGC1α mRNA expression. Patients were divided into high and low PGC1α expression groups based on mean expression. Disease-free survival (y-axis) is plotted against time (x-axis). Differences between groups were assessed using the log-rank test. Hazard ratios (HR) and corresponding 95% confidence intervals (CI) were calculated using a Cox proportional hazards model. UP: upregulated; FDR= false discovery rate; FC=fold change; CAF= cancer associated fibroblast; p.v= p-value. Statistics: Non-parametric t-test (A, H), Fisher exact test (C-D), Log-rank test (J). Error bars indicate s.e.m.

**Supplementary Figure 2. A.** Validation of PGC1α expression by western blot (n=3) in PC3 cell line (PC3 TRIPZ-Pgc1a). **B**. Reactome analysis from differential gene expression upregulated in transcriptomic data comparing PGC1α expressing and not expressing cells. Cut-offs: p-value< 0.05; FC>1.2. **C**. KEGG analysis of differential gene expression upregulated (upper panel) and downregulated (lower panel) in transcriptomic data comparing PGC1α expressing and not expressing cells. Cut-offs: p-value< 0.05; FC> or< 1.2 **D**. 2D cell adhesion to different coatings (collagen I, fibronectin, laminin, vitronectin and tenascin C) of DU145 PGC1α expressing cells (DU145 TRIPZ-Pgc1a). **E**. 3D adhesion to collagen I matrix of DU145 PGC1α expressing cells. **F-G** Doxycycline treatment control on 2D and 3D adhesion respectively using cells with the empty vector used (TRIPZ). Numbers represent p-values. Dox=doxycycline; DEG= differential expression analysis; FC=fold change; IGF= insulin-like growth factor; FA= fatty acid; TCA= tricarboxylic. Statistics: one sample t-test with reference value 1 (D, E, F, G). Numbers represent p-values. Error bars indicate s.e.m.

**Supplementary Figure 3. A**. Validation of PPARGC1A gene expression by RT-qPCR in the tumours generated orthotopically in the prostate (ventral lobe), by injecting PC3 TRIPZ-Pgc1a cells. **B**. E*x vivo b*ioluminescence analysis (total flux measurements) of samples harvested from the brain, inguinal and lumbar lymph nodes (LN, LLN), heart, liver, and kidney by means of total flux measurement. **C**. Volcano plot representing upregulated and downregulated proteins obtained from the LC/MS label-free proteomic analysis of orthotopic prostate tumours. **D**. Table with the top common and consistent candidates whose gene (Supplementary Table S4) and protein (Supplementary Table S5) expression is downregulated upon PGC1α re-expression. **E**. Gene expression analysis of the selected candidates in DU145 TRIPZ and DU145 TRIPZ Pgc1a cells induced with doxycycline (0.5 μg/μL). **F**. Analysis of COL13A1, IL1B and PLAU mRNA in PCa patients stratified according to the mean expression of PGC1α mRNA (PPARGC1A) in TCGA and Taylor datasets. Sample sizes: Taylor database n=131, TCGA provisional n=497. **G**. Analysis of CTHRC1 mRNA expression in epithelial (left panel) and stromal (right panel) compartments of PCa specimens stratified according to the mean expression of PGC1α mRNA. Sample size: Tyekucheva n=25, in which P_high n=10 and P_low n=15. Numbers represent p-values. Dox: doxycycline; FC: fold change; p/s= photons per second; LC/MS= liquid chromatography/mass spectrometry; RNAseq= RNA sequencing; P_high= PPARGC1A high; P_low= PPARGC1A low. Statistics: data was normalized to each no dox condition in E. Non-parametric t-test (A, B, F) and one sample t-test with reference value 1 (E). Numbers represent p-values. Error bars indicate s.e.m.

**Supplementary Figure 4. A**. Analysis of CTHRC1 mRNA expression in PCa patients in Taylor, Varambally, Grasso and Lapointe datasets (Cancertool). **B**. Effect of CTHRC1 silencing on cell proliferation (crystal violet staining) of PC3 and DU145 cell lines. **C-D**. Effect of CTHRC1 silencing in DU145 cells on BrDU incorporation and transwell migration assays (right panel shows representative images of transwell migration). **E**. Effect of CTHRC1 silencing on 2D cell adhesion of DU145 cells to different ECM-coatings (collagen I, fibronectin, laminin and vitronectin). Dox: doxycycline; FC: fold change; sh=short hairpin; Scr=scramble; N=normal; PT= primary tumor; M= Metastasis. Statistics: data was normalized to each no dox condition in B, C, D, E. One sample t-test with reference value 1 (B, C, D, E). Numbers represent p-values. Error bars indicate s.e.m.

**Supplementary Figure 5. A.** Scheme of CTHRC1 re-expression in PGC1α expressing and non-expressing cells. **B-C**. Gene and protein expression validation of CTHRC1 re-expression in PC3 PGC1α expressing and non-expressing cells. **D-E**. Detection of CTHRC1 (ELISA, D;

Western blot, E) in the conditioned media of PGC1α expressing and non-expressing cells with and without CTHRC1 re-expression. **F**. Detection of deglycosylated CTHRC1 recombinant protein by western blot. **G**. Representative imagines of two-photon microscopy of *in ovo* tumors (Figure 5 H). Scale bar represents 20µm and colour coding represents angle degree. Dox: doxycycline; FC: fold change; ND= non-denaturalized; D= denaturalized. Empty h and empty p correspond to control plasmids for hygromycin and puromycin resistance respectively. Statistics: data was normalized to each no dox condition. One sample t-test (B). Numbers represent p-values. Error bars indicate s.e.m.
